# Hydraulic resistance of perivascular spaces in the brain

**DOI:** 10.1101/522409

**Authors:** Jeffrey Tithof, Douglas H. Kelley, Humberto Mestre, Maiken Nedergaard, John H. Thomas

## Abstract

**Background:** Perivascular spaces (PVSs) are annular channels that surround blood vessels and carry cerebrospinal fluid through the brain, sweeping away metabolic waste. *In vivo* observations reveal that they are not concentric, circular annuli, however: the outer boundaries are often oblate, and the blood vessels that form the inner boundaries are often offset from the central axis.

**Methods:** We model PVS cross-sections as circles surrounded by ellipses and vary the radii of the circles, major and minor axes of the ellipses, and two-dimensional eccentricities of the circles with respect to the ellipses. For each shape, we solve the governing Navier-Stokes equation to determine the velocity profile for steady laminar flow and then compute the corresponding hydraulic resistance.

**Results:** We find that the observed shapes of PVSs have lower hydraulic resistance than concentric, circular annuli of the same size, and therefore allow faster, more efficient flow of cerebrospinal fluid. We find that the minimum hydraulic resistance (and therefore maximum flow rate) for a given PVS cross-sectional area occurs when the ellipse is elongated and intersects the circle, dividing the PVS into two lobes, as is common around pial arteries. We also find that if both the inner and outer boundaries are nearly circular, the minimum hydraulic resistance occurs when the eccentricity is large, as is common around penetrating arteries.

**Conclusions:** The concentric circular annulus assumed in recent studies is not a good model of the shape of actual PVSs observed *in vivo*, and it greatly overestimates the hydraulic resistance of the PVS. Our parameterization can be used to incorporate more realistic resistances into hydraulic network models of flow of cerebrospinal fluid in the brain. Our results demonstrate that actual shapes observed *in vivo* are nearly optimal, in the sense of offering the least hydraulic resistance. This optimization may well represent an evolutionary adaptation that maximizes clearance of metabolic waste from the brain.

## Background

It has long been thought that flow of cerebrospinal fluid (CSF) in perivascular spaces plays an important role in the clearance of solutes from the brain [1, 2, 3]. Experiments have shown that tracers injected into the subarachnoid space are transported preferentially into the brain through periarterial spaces at rates much faster than can be explained by diffusion alone [4, 5, 6]. Recent experimental results [7, 8] now show unequivocally that there is pulsatile flow in the perivascular spaces around pial arteries in the mouse brain, with net (bulk) flow in the same direction as the blood flow. These *in vivo* measurements support the hypothesis that this flow is driven primarily by “perivascular pumping” due to motions of the arterial wall synchronized with the cardiac cycle [8]. From the continuity equation (expressing conservation of mass), we know that this net flow must continue in some form through other parts of the system (e.g., along PVSs around penetrating arteries, arterioles, capillaries, venules). The *in vivo* experimental methods of Mestre *et al.* [8] now enable measurements of the size and shape of the perivascular spaces, the motions of the arterial wall, and the flow velocity field in great detail.

With these *in vivo* measurements, direct simulations can in principle predict the observed fluid flow by solving the Navier-Stokes (momentum) equation. A handful of numerical [9, 10, 11, 12, 13] and analytical [14, 15] studies have previously been developed to model CSF flow through PVSs. These studies provide important steps in understanding the fluid dynamics of the entire glymphatic system [3, 16], not only in mice but in mammals generally. However, these studies have been based on idealized assumptions and have typically simulated fluid transport through only a small portion of the brain. Development of a fully-resolved fluid-dynamic model that captures CSF transport through the entire brain is beyond current capabilities for two reasons: (i) the very large computational cost of such a simulation, and (ii) the lack of detailed knowledge of the configuration and mechanical properties of the various flow channels throughout the glymphatic pathway, especially deep within the brain. We note that these limitations and the modest number of publications modeling CSF transport through the brain are in contrast with the much more extensive body of research modeling CSF flow in the spinal canal, which has pursued modeling based on idealized [17, 18, 19], patient-specific [20, 21], and *in vitro* [22] geometries (see the recent review articles [23, 24, 25]).

To simulate CSF transport at a brain-wide scale, a tractable first step is to model the flow using a hydraulic network by estimating the hydraulic resistance of the channels that carry the CSF, starting with the PVSs. This article is restricted to modeling of CSF flow through PVSs in the brain and does not address the question of flow through the brain parenchyma [26, 27], a region where bulk flow phenomena have not been characterized in the same detail as in the PVS. A steady laminar (Poiseuille) flow of fluid down a channel is characterized by a volume flow rate 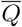 that is proportional to the pressure drop ∆*p* along the channel. The inverse of that proportionality constant is the hydraulic resistance 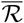. Higher hydraulic resistance impedes flow, such that fewer mL of CSF are pumped per second by a given pressure drop ∆*p*; lower hydraulic resistance promotes flow. Hydraulic resistance is analogous to electrical resistance, which impedes the electrical current driven by a given voltage drop. The hydraulic resistance of a channel for laminar flow can be calculated from the viscosity of the fluid and the length, shape, and cross-sectional area of the channel. We note that prior numerical studies have computed the hydraulic resistance of CSF flow in the spinal canal [28, 29], and a few hydraulic-network models of perivascular flows have been presented, using a concentric circular-annulus configuration of the PVS cross-section (e.g., [12, 30, 31]). As we demonstrate below, the concentric circular annulus is generally not a good model of the cross-section of a PVS. Here we propose a simple but more realistic model that is adjustable and able to approximate the cross-sections of PVSs actually observed in the brain. We then calculate the velocity profile, volume flow rate, and hydraulic resistance for Poiseuille flow with these cross-sections and demonstrate that the shapes of PVSs around pial arteries are nearly optimal.

## Methods

### The basic geometric model of the PVS

In order to estimate the hydraulic resistance of PVSs, we need to know the various sizes and shapes of these spaces *in vivo*. Recent measurements of periarterial flows in the mouse brain by Mestre *et al.* [8] show that the perivascular space (PVS) around the pial arteries is much larger than previously estimated—comparable to the diameter of the artery itself. *In vivo* experiments using fluorescent dyes show similar results [32]. The size of the PVS is substantially larger than that shown in previous electron microscope measurements of fixed tissue. Mestre *et al.* demonstrate that the PVS collapses during fixation: they find that the ratio of the cross-sectional area of the PVS to that of the artery itself is on average about 1.4 *in vivo*, whereas after fixation this ratio is only about 0.14.

The *in vivo* observation of the large size of the PVS around pial arteries is important for hydraulic models because the hydraulic resistance depends strongly on the size of the channel cross-section. For a concentric circular annulus of inner and outer radii *r*_1_ and *r*_2_, respectively, for fixed *r*_1_ the hydraulic resistance scales roughly as (*r*_2_*/r*_1_)^*−*4^, and hence is greatly reduced in a wider annulus. As we demonstrate below, accounting for the actual shapes and eccentricities of the PVSs will further reduce the resistance of hydraulic models.

Figure 1 shows images of several different cross-sections of arteries and surrounding PVSs in the brain, measured *in vivo* using fluorescent dyes [8, 6, 32, 33] or optical coherence tomography [7]. The PVS around a pial artery generally forms an annular region, elongated in the direction along the skull. For an artery that penetrates into the parenchyma, the PVS is less elongated, assuming a more circular shape, but not necessarily concentric with the artery. Note that similar geometric models have been used to model CSF flow in the cavity (ellipse) around the spinal cord (circle) [17, 18].

**Fig. 1:**
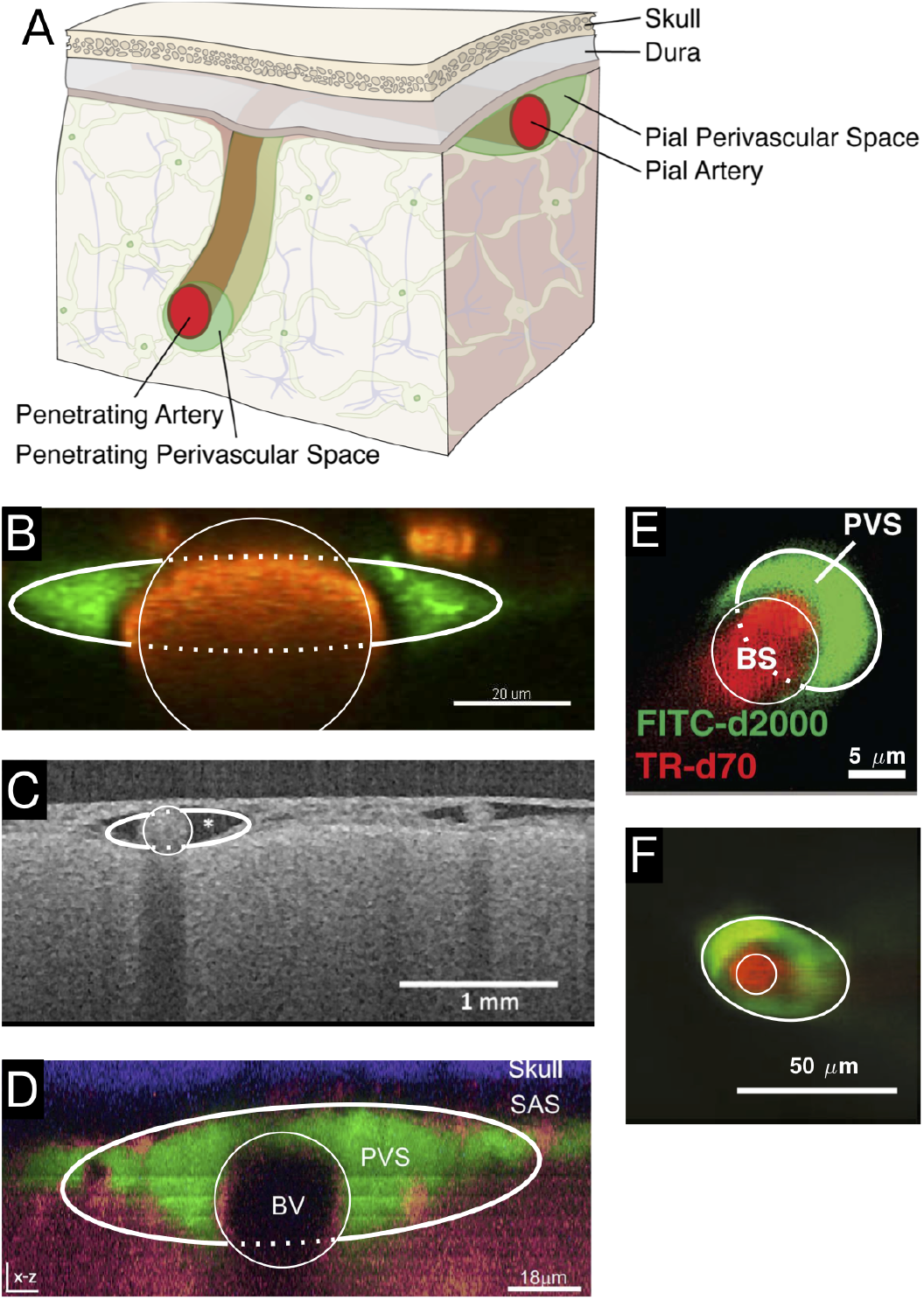
Cross-sections of PVSs from *in vivo* dye experiments. **A** We consider PVSs in two regions: those adjacent to pial arteries and those adjacent to penetrating arteries. **B** PVS surrounding a murine pial artery, adapted from [8]. **C** PVS surrounding a human pial artery, adapted from [7]. **D** PVS surrounding a murine pial artery, adapted from [32]. **E** PVS surrounding a murine descending artery, adapted from [6]. **F** PVS surrounding a murine descending artery, adapted from [33]. For each image B-F, the best-fit inner circular and outer elliptical boundaries are plotted (thin and thick curves, respectively). The model PVS cross-section is the space within the ellipse but outside the circle. The dotted line does not represent an anatomical structure but is included to clearly indicate the fit. The parameter values for these fits are given in Table 1. PVSs surrounding pial arteries are oblate, not circular; PVSs surrounding descending arteries are more nearly circular, but are not concentric with the artery.

We need a simple working model of the configuration of a PVS that is adjustable so that it can be fit to the various shapes that are actually observed, or at least assumed. Here we propose the model shown in Figure 2. This model consists of an annular channel whose cross-section is bounded by an inner circle, representing the outer wall of the artery, and an outer ellipse, representing the outer wall of the PVS. The radius *r*_1_ of the circular artery and the semi-major axis *r*_2_ (*x*-direction) and semi-minor axis *r*_3_ (*y*-direction) of the ellipse can be varied to produce different cross-sectional shapes of the PVS. With *r*_2_ = *r*_3_ *> r*_1_, we have a circular annulus. Generally, for a pial artery, we have *r*_2_ *> r*_3_ *≈ r*_1_: the PVS is annular but elongated in the direction along the skull. For *r*_3_ = *r*_1_ *< r*_2_, the ellipse is tangent to the circle at the top and bottom, and for *r*_3_ *≤ r*_1_ *< r*_2_ the PVS is split into two disconnected regions, one on either side of the artery, a configuration that we often observe for a pial artery in our experiments. We also allow for eccentricity in this model, allowing the circle and ellipse to be non-concentric, as shown in Figure 2B. The center of the ellipse is displaced from the center of the circle by distances *c* and *d* in the *x* and *y* directions, respectively. The model is thus able to match quite well the various observed shapes of PVSs. To illustrate this, in Figure 1 we have drawn the inner and outer boundaries (thin and thick white curves, respectively) of the geometric model that gives a close fit to the actual configuration of the PVS. Specifically, the circles and ellipses plotted have the same centroids and the same normalized second central moments as the dyed regions in the images. We have drawn the full ellipse indicating the outer boundary of the PVS to clearly indicate the fit, but the portion which passes through the artery is plotted with a dotted line to indicate that this does not represent an anatomical structure.

**Table 1:** Geometry and resistance of perivascular spaces visualized *in vivo*. Labels correspond to panel labels in Figure 1. The last column gives the ratio of the hydraulic resistance 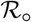 of a circular annulus with the same area ratio *K* to the value 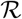 computed for the specified geometry.

| Label | $r_1$ | $r_2$ | $r_3$ | $A_{art}$ | $A_{pvs}$ | $c$ | $d$ |
| --- | --- | --- | --- | --- | --- | --- | --- |
| B | 19.92 $\mu\text{m}$ | 42.1 $\mu\text{m}$ | 8.09 $\mu\text{m}$ | 1169 $\mu\text{m}^2$ | 1059 $\mu\text{m}^2$ | -0.0428 $\mu\text{m}$ | 5.23 $\mu\text{m}$ |
| C | 152.9 $\mu\text{m}$ | 449 $\mu\text{m}$ | 113.7 $\mu\text{m}$ | $6.63 \times 10^4 \mu\text{m}^2$ | $1.577 \times 10^5 \mu\text{m}^2$ | -67.6 $\mu\text{m}$ | 14.84 $\mu\text{m}$ |
| D | 16.53 $\mu\text{m}$ | 58.6 $\mu\text{m}$ | 16.67 $\mu\text{m}$ | 742 $\mu\text{m}^2$ | 2670 $\mu\text{m}^2$ | -4.18 $\mu\text{m}$ | 6.55 $\mu\text{m}$ |
| E | 4.63 $\mu\text{m}$ | 6.83 $\mu\text{m}$ | 5.42 $\mu\text{m}$ | 59.2 $\mu\text{m}^2$ | 113.5 $\mu\text{m}^2$ | -0.513 $\mu\text{m}$ | -4.61 $\mu\text{m}$ |
| F | 7.21 $\mu\text{m}$ | 23.3 $\mu\text{m}$ | 15.40 $\mu\text{m}$ | 155.0 $\mu\text{m}^2$ | 1120 $\mu\text{m}^2$ | 0.1192 $\mu\text{m}$ | -5.74 $\mu\text{m}$ |
| Label | $\alpha$ | $\beta$ | $K$ | $\epsilon_x$ | $\epsilon_y$ | $r_1^4 \mathcal{R} / \mu$ | $\mathcal{R}_o / \mathcal{R}$ |
| B | 2.11 | 0.406 | 0.388 | -0.00215 | 0.263 | 48.0 | 6.45 |
| C | 2.94 | 0.744 | 1.36 | -0.442 | 0.0971 | 3.56 | 2.75 |
| D | 3.54 | 1.008 | 2.71 | -0.253 | 0.396 | 1.01 | 1.62 |
| E | 1.476 | 1.172 | 1.18 | -0.1109 | -0.997 | 3.30 | 4.29 |
| F | 3.24 | 2.14 | 5.93 | 0.0165 | -0.797 | 0.173 | 1.38 |

**Fig. 2:**
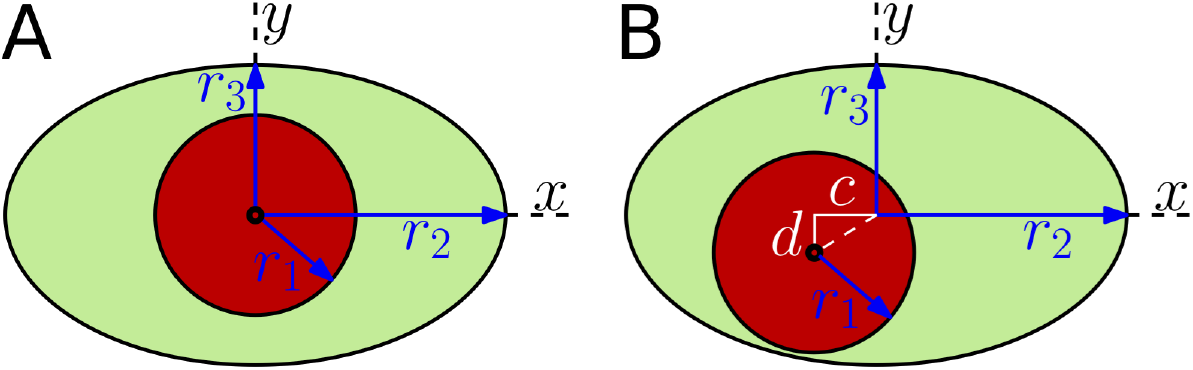
Adjustable geometric models of the cross-section of a PVS, where the circle represents the outer boundary of the artery and the ellipse represents the outer boundary of the PVS. The circle and ellipse may be either **A** concentric or **B** non-concentric. In **A**, the geometry is parameterized by the circle radius *r*_1_ and the two axes of the ellipse *r*_2_ and *r*_3_. In **B**, there are two additional parameters: eccentricities *c* along the *x*-direction and *d* along the *y*-direction.

### Steady laminar flow in the annular tube

We wish to find the velocity distribution for steady, fully developed, laminar viscous flow in our model tube, driven by a uniform pressure gradient in the axial (*z*) direction. The velocity *u*(*x, y*) is purely in the *z*-direction and the nonlinear term in the Navier-Stokes equation is identically zero. The basic partial differential equation to be solved is the *z*-component of the Navier-Stokes equation, which reduces to

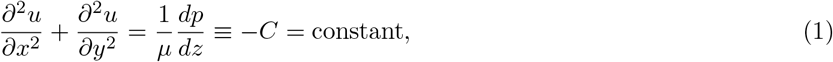

where *µ* is the dynamic viscosity of the CSF. (Note that the pressure gradient *dp/dz* is constant and negative, so the constant *C* we have defined here is positive.) If we introduce the nondimensional variables

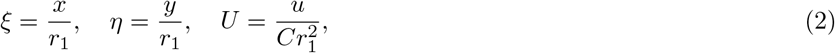

then equation (1) becomes the nondimensional Poisson’s equation

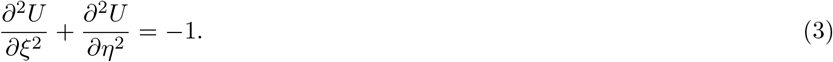

We want to solve this equation subject to the Dirichlet (no-slip) condition *U* = 0 on the inner (circle) and outer (ellipse) boundaries. Analytic solutions are known for simple geometries, and we can calculate numerical solutions for a wide variety of geometries, as described below.

Let *A*_*pvs*_ and *A*_*art*_ denote the cross-sectional areas of the PVS and the artery, respectively. Now, define the nondimensional parameters

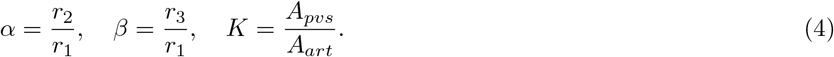

(Note that *K* is also equal to the volume ratio *V_pvs_/V_art_* of a fixed length of our tube model.) When *r*_1_, *r*_2_, *r*_3_, *c*, and *d* have values such that the ellipse surrounds the circle without intersecting it, the cross-sectional areas of the PVS and the artery are given simply by

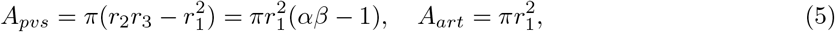

and the area ratio is

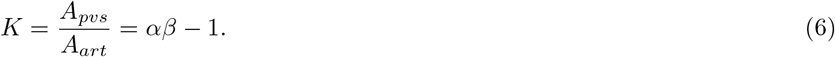

In cases where the ellipse intersects the circle, the determination of *A*_*pvs*_ is more complicated: in this case, equations (5) and (6) are no longer valid, and instead we compute *A*_*pvs*_ numerically, as described in more detail below.

For our computations of velocity profiles in cases with no eccentricity (*c* = *d* = 0), we can choose a value of the area ratio *K*, which fixes the volume of fluid in the PVS, and then vary *α* to change the shape of the ellipse. Thus we generate a two-parameter family of solutions: the value of *β* is fixed by the values of *K* and *α*. In cases where the circle does not protrude past the boundary of the ellipse, the third parameter *β* varies according to *β* = (*K* + 1)*/α*. For *α* = 1 the ellipse and circle are tangent at *x* = *±r*_2_, *y* = 0 and for *α* = *K* + 1 they are tangent at *x* = 0, *y* = *±r*_3_. Hence, for fixed *K*, the circle does not protrude beyond the ellipse for *α* in the range 1 *≤ α ≤ K* + 1. For values of *α* outside this range, we have a two-lobed PVS, and the relationship among *K*, *α*, and *β* is more complicated.

The dimensional volume flow rate 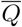 is found by integrating the velocity-profile

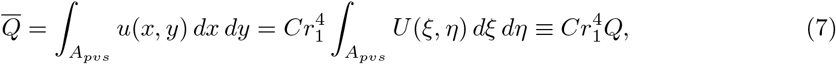

where 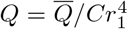 is the dimensionless volume flow rate. The hydraulic resistance 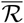 is given by the relation 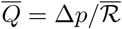, where ∆*p* = (*−dp/dz*)*L* is the pressure drop over a length *L* of the tube. For our purposes, it is better to define a hydraulic resistance *per unit length*, 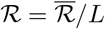, such that

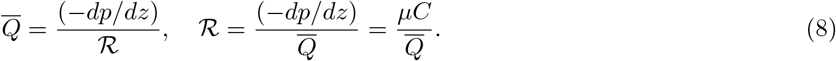

We can use computed values of *Q* to obtain values of the hydraulic resistance 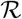. From equations (7) and (8), we have

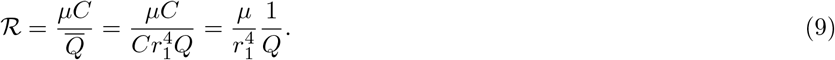

We can then plot the scaled, dimensionless resistance 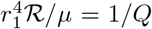 as a function of (*α − β*)*/K* (shape of the ellipse) for different values of *K* (area ratio).

For viscous flows in ducts of various cross-sections, the hydraulic resistance is often scaled using the *hydraulic radius r*_h_ = 2*A/P*, where *A* is the cross-sectional area of the duct and *P* is the wetted perimeter. In the case of our annular model, however, the hydraulic radius *r*_h_ = 2*A_pvs_/P* is not a useful quantity: when the inner circle lies entirely within the outer ellipse, both *A*_*pvs*_ and *P*, and hence *r*_h_, are independent of the eccentricity, but (as shown below) the hydraulic resistance varies with eccentricity.

### Numerical methods

In order to solve Poisson’s equation (3) subject to the Dirichlet condition *U* = 0 on the inner and outer boundaries of the PVS, we employ the Partial Differential Equation (PDE) Toolbox in MATLAB. This PDE solver utilizes finite-element methods and can solve Poisson’s equation in only a few steps. First, the geometry is constructed by specifying a circle and an ellipse (the ellipse is approximated using a polygon with a high number of vertices, typically 100). Eccentricity may be included by shifting the centers of the circle and ellipse relative to each other. We specify that the equation is to be solved in the PVS domain corresponding to the part of the ellipse that does not overlap with the circle. We next specify the Dirichlet boundary condition *U* = 0 along the boundary of the PVS domain and the coefficients that define the nondimensional Poisson’s equation (3). Finally, we generate a fine mesh throughout the PVS domain, with a maximum element size of 0.02 (nondimensionalized by *r*_1_), and MATLAB computes the solution to equation (3) at each mesh point. The volume flow rate is obtained by numerically integrating the velocity profile over the domain. Choosing the maximum element size of 0.02 ensures that the numerical results are converged. Specifically, we compare the numerically obtained value of the flow rate *Q* for a circular annulus to the analytical values given by equation (11) or equation (12) below to ensure that the numerical results are accurate to within 1%.

For the case where the circle protrudes beyond the boundary of the ellipse, equations (5) and (6) do not apply. We check for this case numerically by testing whether any points defining the boundary of the circle extrude beyond the boundary of the ellipse. If so, we compute the area ratio *K* numerically by integrating the area of the finite elements in the PVS domain (*A*_*art*_ is known but *A*_*pvs*_ is not). In cases where we want to fix *K* and vary the shape of the ellipse (e.g. Fig. 5A), it is necessary to change the shape of the ellipse iteratively until *K* converges to the desired value. We do so by choosing *α* and varying *β* until *K* converges to its desired value within 0.01%.

### Analytical solutions

There are two special cases for which there are explicit analytical solutions, and we can use these solutions as checks on the numerical method.

#### The concentric circular annulus

For a concentric circular annulus we have *c* = *d* = 0, *r*_2_ = *r*_3_ *> r*_1_, *α* = *β >* 1, and *K* = *α*^2^ *−* 1. Let *r* be the radial coordinate, and *ρ* = *r/r*_1_ be the corresponding dimensionless radial coordinate. The dimensionless velocity profile is axisymmetric, and is given by White [34], p. 114:

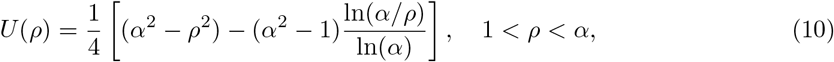

and the corresponding dimensionless volume flux rate is given by:

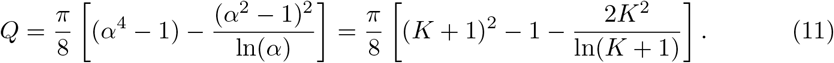

#### The eccentric circular annulus

There is also an analytical solution for the case of an eccentric circular annulus, in which the centers of the two circles do not coincide [34, 35]. Let *c* denote the radial distance between the two centers. Then, in cases where the two circles do not intersect, the dimensionless volume flow rate is given by White [34], p. 114:

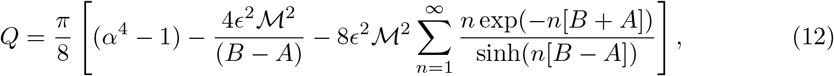

where *ε* = *c/r*_1_ is the dimensionless eccentricity and

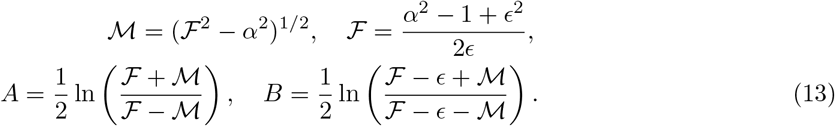

From this solution, it can be shown that increasing the eccentricity substantially increases the flow rate (see Figs. 3–10 in [34]). This solution can be used as a check on the computations of the effect of eccentricity in our model PVS in the particular case where the outer boundary is a circle.

**Fig. 3:**
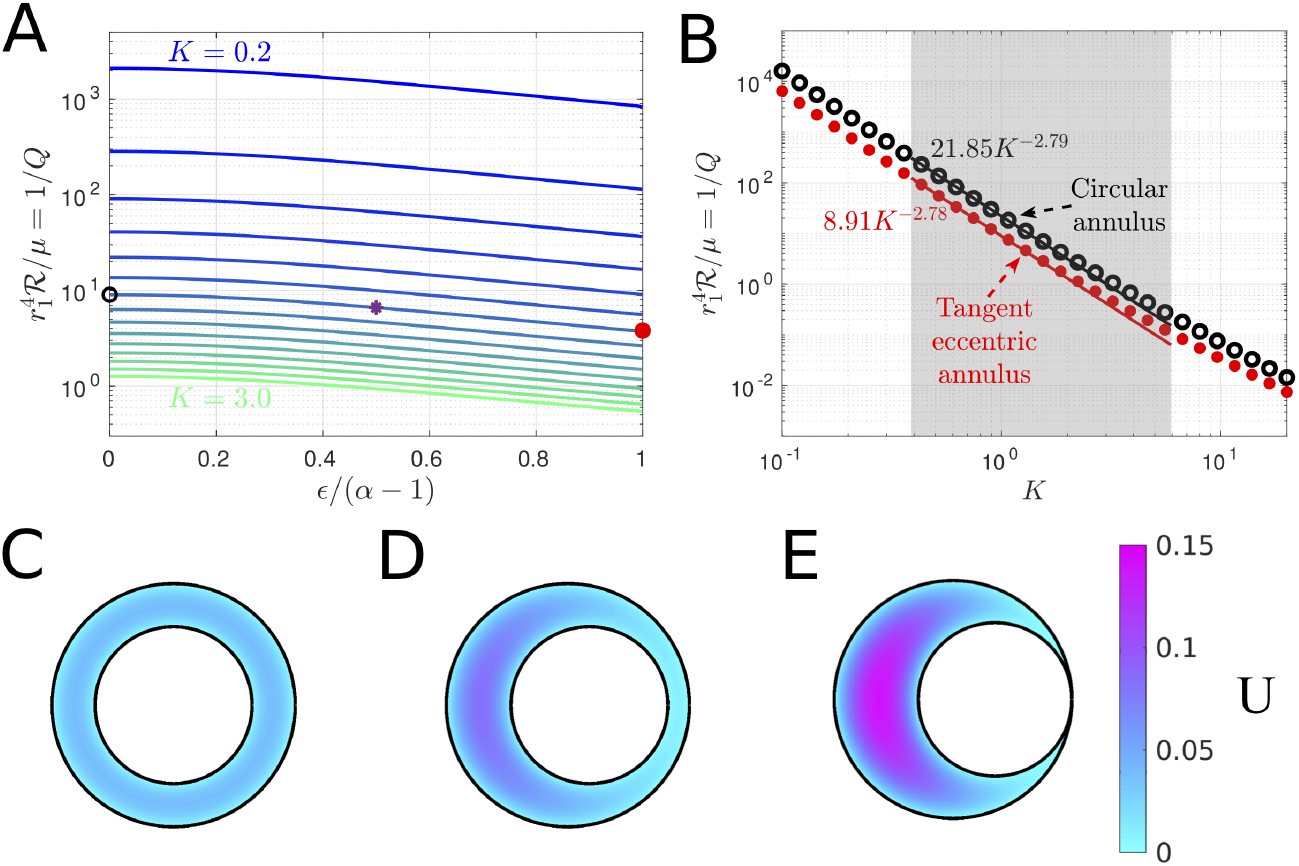
Hydraulic resistance and velocity profiles in eccentric circular annuli modeling PVSs surrounding penetrating arteries. **A** Plots of hydraulic resistance 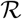 for an eccentric circular annulus, as a function of the relative eccentricity ε/(α − 1), for various fixed values of the area ratio *K* = α^2^ − 1 ranging in steps of 0.2, computed using equation (12). **B** Plots of the hydraulic resistance (red dots) for the tangent eccentric circular annulus (defined as ε/(α − 1) = 1) as a function of the area ratio *K*. Also plotted, for comparison, is the hydraulic resistance of the concentric circular annulus for each value of *K*. The shaded region indicates the range of *K* observed *in vivo* for PVSs. Power laws are indicated that fit the points well through most of the shaded region. C-E Velocity profiles for three different eccentric circular annuli with increasing eccentricity (with K = 1.4 held constant): (C) ε = 0 (concentric circular annulus), (D) ε = 0.27 (eccentric circular annulus), and (E) ε = 0.55 (tangent eccentric circular annulus). The black circle, purple asterisk, and red dot in A indicate the hydraulic resistance of the shapes shown in C-E, respectively. The volume flow rates for the numerically calculated profiles shown in C-E agree with the analytical values to within 0.3%. As eccentricity increases hydraulic resistance decreases and volume flow rate increases.

## Results

### The eccentric circular annulus

The eccentric circular annulus is a good model for the PVSs around some penetrating arteries (see Fig. 1E,F), so it is useful to show how the volume flow rate and hydraulic resistance vary for this model. This is done in Figure 3A, where the hydraulic resistance (inverse of the volume flow rate) is plotted as a function of the dimensionless eccentricity *c/*(*r*_2_ *− r*_1_) = *ε/*(*α −* 1) for various values of the area ratio *K* = *α*^2^ *−* 1. The first thing to notice in this plot is how strongly the hydraulic resistance depends on the cross-sectional area of the PVS (i.e., on *K*). For example, in the case of a concentric circular annulus (*ε* = 0), the resistance decreases by about a factor of 1700 as the area increases by a factor of 15 (*K* goes from 0.2 to 3.0).

For fixed *K*, the hydraulic resistance decreases monotonically with increasing eccentricity (see Fig. 3A). This occurs because the fluid flow concentrates more and more into the wide part of the gap, where it is farther from the walls and thus achieves a higher velocity for a given shear stress (which is fixed by the pressure gradient). (This phenomenon is well known in hydraulics, where needle valves tend to leak badly if the needle is flexible enough to be able to bend to one side of the circular orifice.) The increase of flow rate (decrease of resistance) is well illustrated in Figures 3C–E, which show numerically computed velocity profiles (as color maps) at three different eccentricities. We refer to the case where the inner circle touches the outer circle (*ε/*(*α −* 1) = 1) as the “tangent eccentric circular annulus.”

We have plotted the hydraulic resistance as a function of the area ratio *K* for the concentric circular annulus and the tangent eccentric circular annulus in Figure 3B. This plot reveals that across a wide range of area ratios, the tangent eccentric circular annulus (shown in Fig. 3E) has a hydraulic resistance that is approximately 2.5 times lower than the concentric circular annulus (shown in Fig. 3C), for a fixed value of *K*. Intermediate values of eccentricity (0 *≤ ε/*(*α −* 1) *≤* 1), where the inner circle does not touch the outer circle (e.g., Fig. 3D) correspond to a reduction in hydraulic resistance that is less than a factor of 2.5. The variation with *K* of hydraulic resistance of the tangent eccentric annulus fits reasonably well to a power law 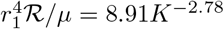 throughout most of the range of observed *K* values, indicated by the gray shaded region in Figure 3B.

### The concentric elliptical annulus

Now we turn to the results for the elliptical annulus in the case where the ellipse and the inner circle are concentric. Figure 4 shows numerically computed velocity profiles for three different configurations with the same area ratio (*K* = 1.4): a moderately elongated annulus, the case where the ellipse is tangent to the circle at the top and bottom, and a case with two distinct lobes. A comparison of these three cases with the concentric circular annulus (Fig. 3B) shows quite clearly how the flow is enhanced when the outer ellipse is flattened, leading to spaces on either side of the artery with wide gaps in which much of the fluid is far from the boundaries and the shear is reduced. However, Figure 4C shows a reduction in the volume flow rate (i.e. less pink in the velocity profile) compared to Figures 4A,B, showing that elongating the outer ellipse too much makes the gaps narrow again, reducing the volume flow rate (increasing the hydraulic resistance). This results suggests that, for a given value of *K* (given cross-sectional area), there is an optimal value of the elongation *α* that maximizes the volume flow rate (minimizes the hydraulic resistance).

**Fig. 4:**
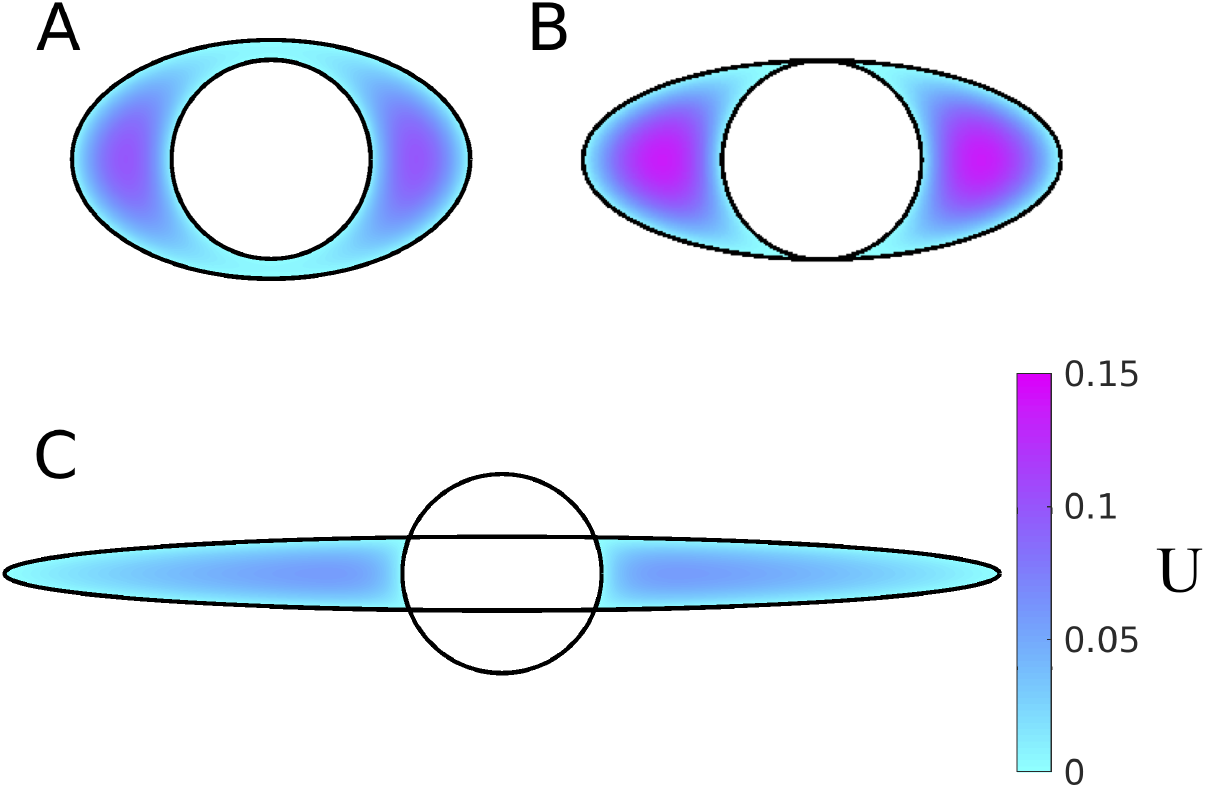
Example velocity profiles in concentric elliptical annuli modeling PVSs surrounding pial arteries. The color maps show velocity profiles for three different shapes of the PVS, all with *K* = 1.4: **A** open PVS (α = 2, β = 1.2), **B** ellipse just touching circle (α = 2.4, β = 1), and **C** two-lobe annulus (α = 5, β = 0.37). Hydraulic resistance is lowest and flow is fastest for intermediate elongation, suggesting the existence of optimal shape that maximizes flow.

To test this hypothesis, we computed the volume flow rate and hydraulic resistance as a function of the shape parameter (*α − β*)*/K* for several values of the area ratio *K*. The results are plotted in Figure 5A. Note that the plot is only shown for (*α −β*)*/K ≥* 0, since the curves are symmetric about (*α −β*)*/K* = 0. The left end of each curve ((*α − β*)*/K* = 0) corresponds to a circular annulus, and the black circles indicate the value of 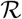 given by the analytical solution in equation (11). These values agree with the corresponding numerical solution to within 1%. The resistance varies smoothly as the outer elliptical boundary becomes more elongated, and our hypothesis is confirmed: for each curve, the hydraulic resistance reaches a minimum value at a value of (*α −β*)*/K* that varies with *K*, such that the corresponding shape is optimal for fast, efficient CSF flow. Typically, the resistance drops by at least a factor of two as the outer boundary goes from circular to the tangent ellipse. If we elongate the ellipse even further (beyond the tangent case), thus dividing the PVS into two separate lobes, the resistance continues to decrease but reaches a minimum and then increases. The reason for this increase is that, as the ellipse becomes highly elongated, it forms a narrow gap itself, and the relevant length scale for the shear in velocity is the width of the ellipse, not the distance to the inner circle. For small values of *K*, we find that the optimal shape parameter (*α − β*)*/K* tends to be large and the ellipse is highly elongated, while for large values of *K* the optimal shape parameter is small. The velocity profiles for three optimal configurations (for *K* = 0.4, 1.4, and 2.4) are plotted in Figures 5C–E.

**Fig. 5:**
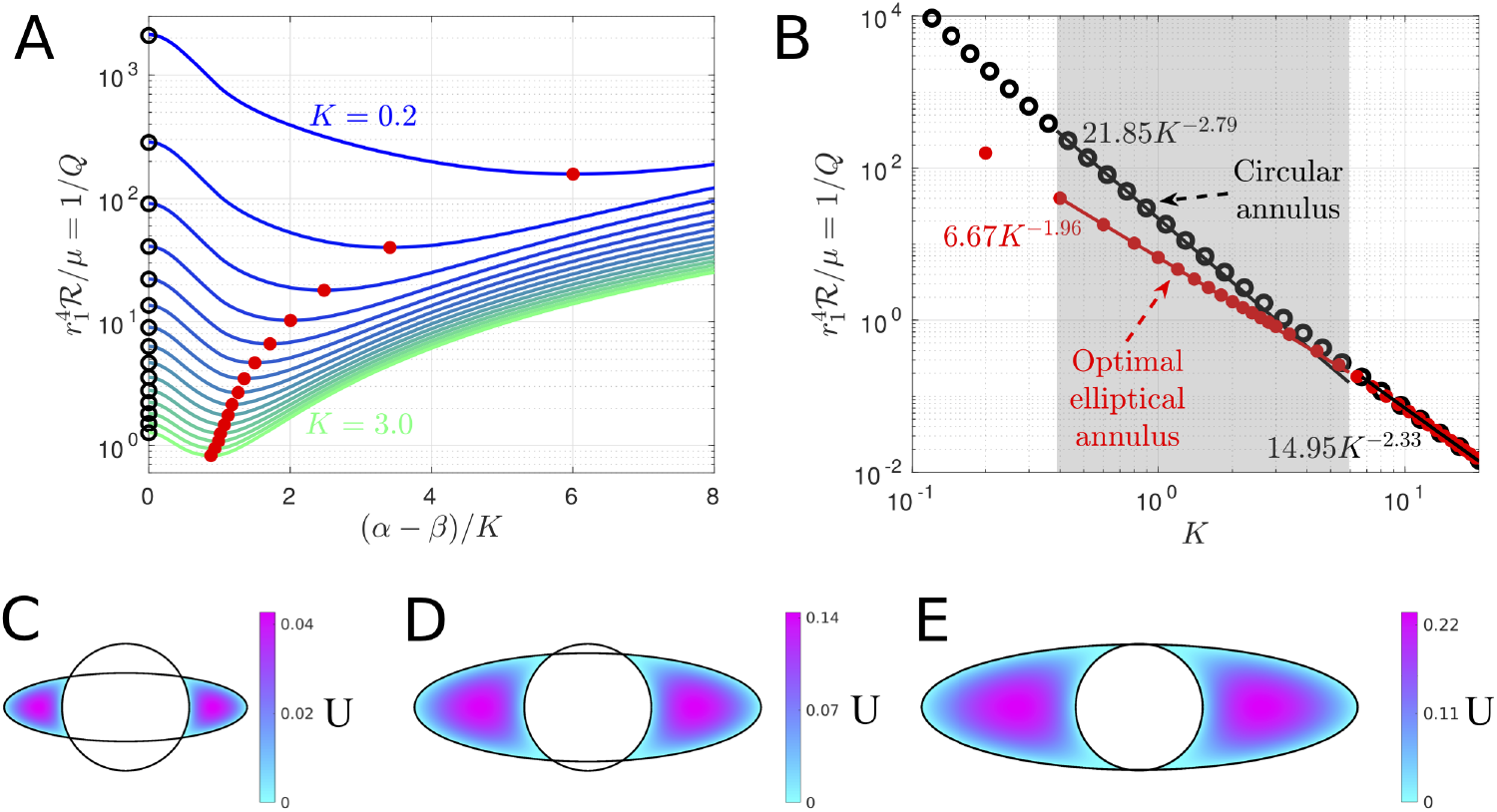
Hydraulic resistance of concentric elliptical annuli modeling PVSs surrounding pial arteries. **A** Hydraulic resistance 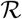 as a function of (α − β)/*K* for various fixed values of the area ratio *K* ranging in steps of 0.2. The black circles indicate the analytic value for the circular annulus, provided by equation (11). Red dots indicate optimal shapes, which have minimum 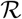 for each fixed value of *K*. **B** Plots of the hydraulic resistance (red dots) for the optimal concentric elliptical annulus as a function of the area ratio *K*. Also plotted, for comparison, is the hydraulic resistance of the concentric circular annulus for each value of *K*. The shaded region indicates the range of *K* observed *in vivo* for PVSs. The two curves in the shaded region are well represented by the power laws shown. For larger values of *K* (larger than actual PVSs) the influence of the inner boundary becomes less significant and the curves converge to a single power law. **C-E** Velocity profiles for the optimal shapes resulting in the lowest hydraulic resistance, with fixed *K* = 0.4, 1.4, and 2.4, respectively. The optimal shapes look very similar to the PVSs surrounding pial arteries (Fig. 1B-D).

The hydraulic resistance of shapes with optimal elongation also varies with the area ratio *K*, as shown in Figure 5B. As discussed above, the resistance decreases rapidly as *K* increases and is lower than the resistance of concentric, circular annuli, which are also shown. We find that the optimal elliptical annulus, compared to the concentric circular annulus, provides the greatest reduction in hydraulic resistance for the smallest area ratios *K*. Although the two curves converge as *K* grows, they differ substantially throughout most of the range of normalized PVS areas observed *in vivo*. We find that the variation with *K* of hydraulic resistance of optimal shapes fits closely to a power law 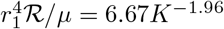.

### The eccentric elliptical annulus

We have also calculated the hydraulic resistance for cases where the outer boundary is elliptical and the inner and outer boundaries are not concentric (see Fig. 2B). For this purpose, we introduce the nondimensional eccentricities

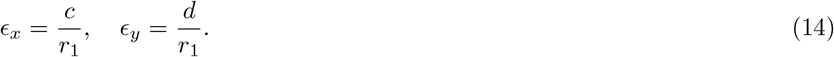

The hydraulic resistance is plotted in Figures 6A,B as a function of *ε_x_* and *ε_y_*, respectively, and clearly demonstrates that adding any eccentricity decreases the hydraulic resistance, similar to the eccentric circular annulus shown in Figure 3. In the case where the outer boundary is a circle 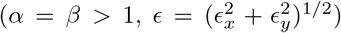 we employ the analytical solution (12) as a check on the numerical solution: they agree to within 0.4%. Two example velocity profiles are plotted in Figures 6C,D. Comparing these profiles to the concentric profile plotted in Figure 4A clearly shows that eccentricity increases the volume flow rate (decreases the hydraulic resistance).

**Fig. 6:**
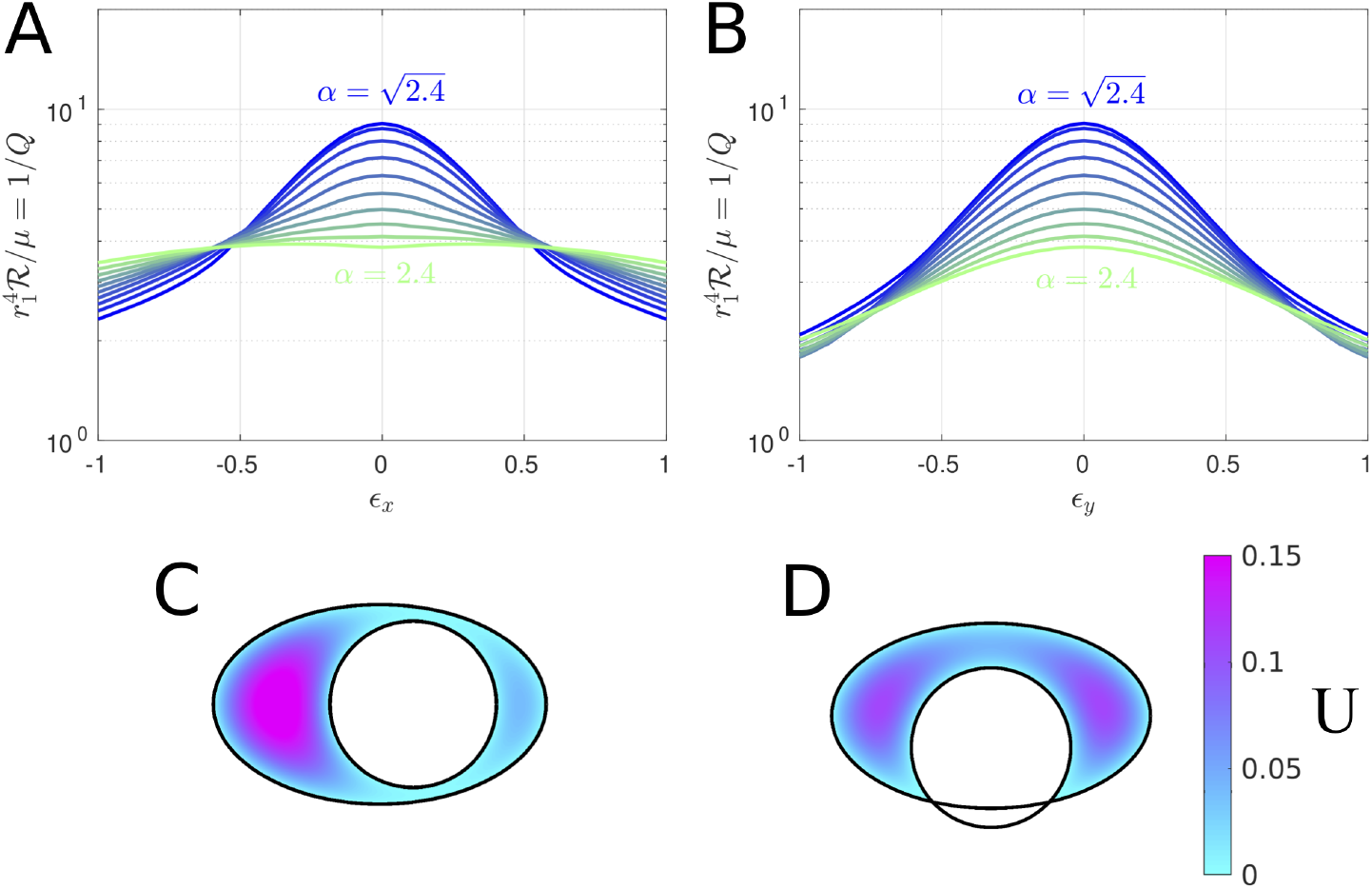
The effects of eccentricity on hydraulic resistance of elliptical annuli modeling PVSs surrounding pial arteries. Hydraulic resistance 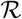 as a function of **A** ε_x_ or **B** ε_y_ for several values of α. Color maps of the velocity profiles for **C** α = 2, ε_x_ = 0.4, ε_y_ = 0 and **D** α = 2, ε_x_ = 0, ε_y_ = −0.4. *K* = 1.4 for all plots shown here. Circular annuli have 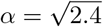, and annuli with 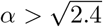 have *r*_2_ > *r*_3_. For a fixed value of α, any nonzero eccentricity increases the flow rate and reduces the hydraulic resistance.

### In vivo PVSs near pial arteries are nearly optimal in shape

We can compute the velocity profiles for the geometries corresponding to the actual pial PVSs shown in Figures 1B–D (dotted and solid white lines). The parameters corresponding to these fits are provided in Table 1 and are based on the model shown in Figure 2B, which allows for eccentricity. Figure 7A shows how hydraulic resistance varies with elongation for non-concentric PVSs having the same area ratio *K* and eccentricities *ε_x_* and *ε_y_* as the ones in Figures 1B–D. The computed values of the hydraulic resistance of the actual observed shapes are plotted as purple triangles. For comparison, velocity profiles for the optimal elongation and the exact fits provided in Table 1 are shown in Figure 7B-D. Clearly the hydraulic resistances of the shapes observed *in vivo* are very close to the optimal values, but systematically shifted to slightly more elongated shapes. Even when (*α − β*)*/K* differs substantially between the observed shapes and the optimal ones, the hydraulic resistance 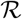, which sets the pumping efficiency and is therefore the biologically important parameter, matches the optimal value quite closely.

**Fig. 7:**
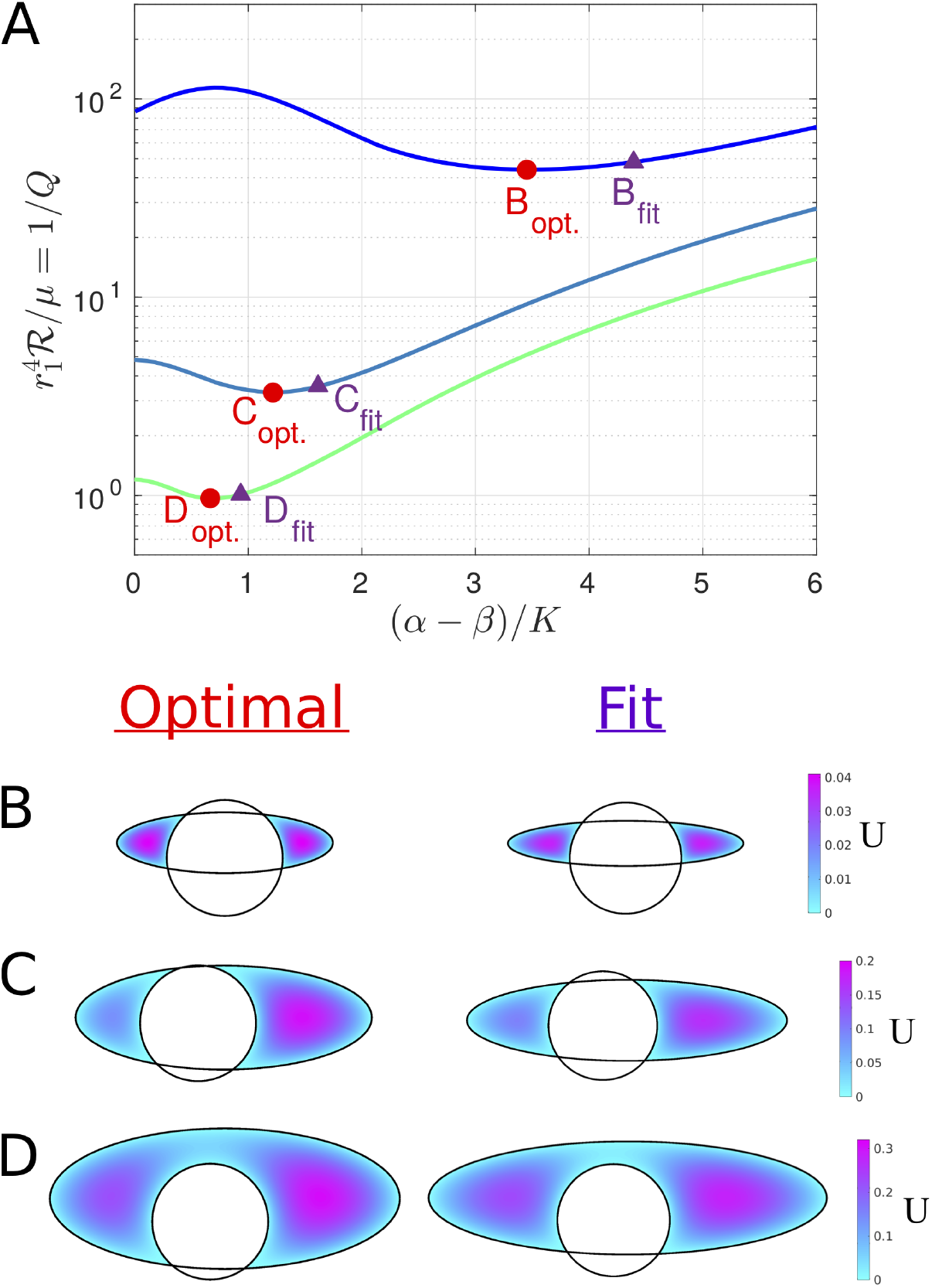
Actual PVS cross-sections measured *in vivo* are nearly optimal. **A** Hydraulic resistance 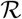 as a function of (α − β)/*K* in which α varies and the values of the area ratio *K* and eccentricities ε_x_ and ε_y_ are fixed corresponding to the fitted values obtained in Table 1. Values corresponding to plots B-D are indicated. **B-D** Velocity profiles for the optimal value of α (left column), which correspond to the minimum value of 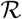 on each curve in **A**, and velocity profiles for the exact fit provided in Table 1 (right column) and plotted in Fig. 1B-D, respectively. The shape of the PVS measured *in vivo* is nearly optimal.

## Discussion

In order to understand the glymphatic system, and various effects on its operation, it will be very helpful to develop a predictive hydraulic model of CSF flow in the PVSs. Such a model must take into account two important recent findings: (i) the PVSs, as measured *in vivo*, are generally much larger than the size determined from post-fixation data [7, 8, 32] and hence offer much lower hydraulic resistance; and (ii) (as we demonstrate in this paper) the concentric circular annulus model is not a good geometric representation of an actual PVS, as it overestimates the hydraulic resistance. With these two factors accounted for, we can expect a hydraulic-network model to produce results in accordance with the actual bulk flow now observed directly in particle tracking experiments [7, 8]. The relatively simple, adjustable model of a PVS that we present here can be used as a basis for calculating the hydraulic resistance of a wide range of observed PVS shapes, throughout the brain and spinal cord. Our calculations demonstrate that accounting for PVS shape can reduce the hydraulic resistance by a factor as large as 6.45 (see Table 1).

We raise the intriguing possibility that the non-circular and eccentric configurations of PVSs surrounding pial arteries are an evolutionary adaptation that lowers the hydraulic resistance and permits faster bulk flow of CSF. The *in vivo* images (e.g., those in Fig. 1B–D) reveal that the cross-section of the PVS around a pial artery is not a concentric circular annulus, but instead is significantly flattened and often consists of two separate lobes positioned symmetrically on each side of the artery. Tracers are mostly moving within these separate tunnels and only to a limited extent passing between them. Our imaging of tens of thousands of microspheres has revealed that crossing is rare, indicating almost total separation between the two tunnels. The arrangement of the two PVS lobes surrounding a pial artery not only reduces the hydraulic resistance but may also enhance the stability of the PVS and prevent collapse of the space during excessive movement of the brain within the skull. Additionally, PVSs with wide spaces may facilitate immune response by allowing macrophages to travel through the brain, as suggested by Schain *et al.* [32]. We note that if CSF flowed through a cylindrical vessel separate from the vasculature (not an annulus), hydraulic resistance would be even lower. However, there are reasons that likely require PVSs to be annular and adjacent to the vasculature, including: (i) arterial pulsations drive CSF flow [8], and (ii) astrocyte endfeet, which form the outer boundary of the PVS, regulate molecular transport from both arteries and CSF [36, 37].

The configuration of PVSs surrounding penetrating arteries in the cortex and striatum is largely unknown [38]. To our knowledge, all existing models are based on information obtained using measurements from fixed tissue. Our own impression, based on years of *in vivo* imaging of CSF tracer transport, is that the tracers distribute asymmetrically along the wall of penetrating arteries, suggesting that the PVSs here are eccentric. Clearly, we need new *in vivo* techniques that produce detailed maps of tracer distribution along penetrating arteries. Regional differences may exist, as suggested by the finding that, in the human brain, the striate branches of the middle cerebral artery are surrounded by three layers of fibrous membrane, instead of the two layers that surround cortical penetrating arteries [38]. Accurately characterizing the shapes and sizes of the most distal PVSs along the arterial tree is very important, as prior work [31] suggests the hydraulic resistance is largest there. We speculate that the configuration of the PVSs at these locations may be optimal as well.

An intriguing possibility for future study is that minor changes in the configuration of PVS spaces may contribute to the sleep-wake regulation of the glymphatic system [39]. Also, age-dependent changes of the configuration of PVSs may increase the resistance to fluid flow, possibly contributing to the increased risk of amyloid-beta accumulation associated with aging [40]. Similarly, reactive remodeling of the PVSs in the aftermath of a traumatic brain injury may increase the hydraulic resistance of PVSs and thereby increase amyloid-beta accumulation.

There are limitations to the modeling presented here, which can be overcome by straightforward extensions of the calculations we have presented. We have intentionally chosen a relatively simple geometry in order to show clearly the dependence of the hydraulic resistance on the size, shape, and eccentricity of the PVS. However, the fits presented in Figure 1B–F are imperfect and could be better captured using high-order polygons, which is an easy extension of the numerical method we have employed. Our calculations have been performed assuming that PVSs are open channels, which is arguably justified – at least for PVSs around pial arteries – by the smooth trajectories observed for 1 *µ*m beads flowing through PVSs and the observation that these spaces collapse during the fixation process [8]. However, the implementation of a Darcy-Brinkman model to capture the effect of porosity would simply increase the resistance 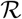, given a fixed flow rate *Q* and Darcy number *Da*, by some multiplicative constant.

The hydraulic resistances we have calculated are for steady laminar flow driven by a constant overall pressure gradient. However, recent quantitative measurements in mice have offered substantial evidence demonstrating that CSF flow in PVSs surrounding the middle cerebral artery is pulsatile, driven by peristaltic pumping due to arterial wall motions generated by the heartbeat, with mean (bulk) flow in the same direction as the blood flow [8]. We hypothesize that this “perivascular pumping” occurs mainly in the periarterial spaces around the proximal sections of the main cerebral arteries: at more distal locations the wall motions become increasingly passive, and the flow is driven mainly by the oscillating pressure gradient generated by the perivascular pumping upstream. Viscous, incompressible duct flows due to oscillating pressure gradients are well understood: it is a linear problem, and analytical solutions are known for a few simple duct shapes. The nature of the solution depends on the *dynamic Reynolds number R_d_* = *ωℓ*^2^*/ν*, where *ω* is the angular frequency of the oscillating pressure gradient, *ν* is the kinematic viscosity, and *ℓ* is the length scale of the duct (e.g, the inner radius of a circular pipe, or the gap width for an annular pipe). (Alternatively, the *Womersley number* 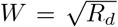 is often used in biofluid mechanics.) When *R_d_ <<* 1, as it is in the case of flows in PVSs,^[1]^ the velocity profile at any instant of time is very nearly that of a steady laminar flow, and the profile varies in time in phase with the oscillating pressure gradient (see White [34], sec. 3-4.2). In this case, the average (bulk) volume flow rate will be inversely proportional to exactly the same hydraulic resistance that applies to steady laminar flow. Hence, the hydraulic resistances we have computed here will apply to PVSs throughout the brain, except for proximal sections of main arteries where the perivascular pumping is actually taking place.

In periarterial spaces where the perivascular pumping is significant, the picture is somewhat different. Here, the flow is actively driven by traveling wave motions of the arterial wall, or in the context of our model PVS, waves along the inner circular boundary. In the case of an elliptical outer boundary, we expect the flow to be three-dimensional, with secondary motions in the azimuthal direction (around the annulus, not down the channel), even though the wave along the inner boundary is axisymmetric. Although we have not yet modeled this flow, we can offer a qualitative description based on an analytical solution for perivascular pumping in the case of concentric circular cylinders [14]. The effectiveness of the pumping scales as (*b/ℓ*)^2^, where *b* is the amplitude of the wall wave and *ℓ* is the width of the gap between the inner and outer boundaries. For the case of a concentric circular annulus, the gap width *ℓ* and hence the pumping effectiveness are axisymmetric, and therefore the resulting flow is also axisymmetric. For an elliptical outer boundary, however, the gap width *ℓ* varies in the azimuthal direction and so will the pumping effectiveness. Hence, there will be pressure variations in the azimuthal direction that will drive a secondary, oscillatory flow in the azimuthal direction, and as a result the flow will be non-axisymmetric and the streamlines will wiggle in the azimuthal direction. Increasing the aspect ratio *r*_2_*/r*_3_ of the ellipse for a fixed area ratio will decrease the flow resistance but will also decrease the overall pumping efficiency, not only because more of the fluid is placed farther from the artery wall, but also, in cases where the PVS is split into two lobes, not all of the artery wall is involved in the pumping. Therefore, we expect that there will be an optimal aspect ratio of the outer ellipse that will produce the maximum mean flow rate due to perivascular pumping, and that this optimal ratio will be somewhat different from that which just produces the lowest hydraulic resistance. We speculate that evolutionary adaptation has produced shapes of actual periarterial spaces around proximal sections of main arteries that are nearly optimal in this sense.

## Conclusions

Perivascular spaces, which are part of the glymphatic system [6], provide a route for rapid influx of cerebrospinal fluid into the brain and a pathway for the removal of metabolic wastes from the brain. In this study, we have introduced an elliptical annulus model that captures the shape of PVSs more accurately than the circular annulus model that has been used in all prior modeling studies. We have demonstrated that for both the circular and elliptical annulus models, non-zero eccentricity (i.e., shifting the inner circular boundary off center) decreases the hydraulic resistance (increases the volume flow rate) for PVSs. By adjusting the shape of the elliptical annulus with fixed PVS area and computing the hydraulic resistance, we found that there is an optimal PVS elongation for which the hydraulic resistance is minimized (the volume flow rate is maximized). We find that these optimal shapes closely resemble actual pial PVSs observed *in vivo*, suggesting such shapes may be a result of evolutionary optimization.

The elliptical annulus model introduced here offers an improvement for future hydraulic network models of the glymphatic system, which may help reconcile the discrepancy between the small PVS flow speeds predicted by many models and the relatively large flow speeds recently measured *in vivo* [7, 8]. Our proposed modeling improvements can be used to obtain simple scaling laws, such as the power laws obtained for the tangent eccentric circular annulus in Figure 3B or the optimal elliptical annulus in Figure 5B.

## Abbreviations

CSF: cerebrospinal fluid
PVS: perivascular space

## Author contributions

JHT developed the theoretical ideas and the geometric model and outlined the calculations. JT and DHK carried out the calculations. HM and MN provided information on actual PVS shapes and flows. JHT, JT, and DHK analyzed the results and wrote the paper.

## Acknowledgements

We thank Dan Xue for assistance with illustrations.

## Competing interests

The authors declare that they have no competing interests.

## Availability of data and materials

All data generated and analyzed in the course of this study are available from the corresponding author upon reasonable request.

## Ethics approval and consent to participate

Not applicable.

## Funding

This work was supported by a grant from the NIH/National Institute of Aging (RF1 AG057575-01 to MN, JHT, and DHK).

## Footnotes

[1] For example, for *ω* = 25.13 s^*−*1^ (corresponding to a pulse rate of 240 bpm), *l*= 20 *µ*m, and *ν* = 7.0 *×* 10^*−*7^m^2^ s^*−*1^, we have *R*_*d*_ = 1.4 *×* 10^*−*2^.

